# Site-specific replacement of large-scale DNA fragments in human cells

**DOI:** 10.1101/2025.09.14.676134

**Authors:** Hojun Jung, Bada Jeong, Yong-Woo Kim, Chanju Jung, Heesoo Uhm, Hyoungrak Kim, Ye Eun Oh, Yoonseo Park, Yeji Lee, Sangsu Bae

## Abstract

Despite advances in genome editing^1-4^, precisely replacing large-scale fragments in human cells remains a significant challenge. Here, we present a site-specific gene replacement tool, named Prime Assembly (PA), which adapts prime editors to produce one or two pairs of 3’-flaps on both the genome and donor plasmids. These 3’-flaps anneal to each other precisely, similar to “Gibson Assembly” in DNA oligonucleotides^5^, allowing megabase-scale genomic excision and/or kilobase-scale donor insertion at the gene of interest. The PA system achieves an efficiency of up to 57.8% in replacing endogenous sequences with a 2.9 kbp donor DNA fragment in HEK293T cells, and presents an accuracy of more than 99% for integrated PA fragments. Ultimately, the site-specific replacement of large-scale coding sequences (CDSs) in disease-related genes can restore gene function across numerous patients with different mutations, providing a gene implantation technique for genome editing and a universal therapeutic approach.

## Main Text

Genetic disorders arise from a wide spectrum of pathogenic mutations scattered across large genomic regions^6,7^. Meanwhile, CRISPR-associated tools enable the direct correction of endogenous gene mutations, creating novel opportunities in therapeutics^1-4,8-13^. Indeed, point mutations can be corrected by base editors (BEs)^8,9^; meanwhile, prime editing (PE) tools can resolve insertion and deletion (indel) mutations as small as several tens of base pairs (bp)^12^. However, the precise correction or replacement of large-scale DNA sequences (>kbp) in human cells currently requires technically challenging strategies, such as restoring short tandem repeats (STRs) and site-specific integration of normal genes. Nonetheless, CRISPR nuclease-based gene correction methods, including homology-directed repair (HDR)^3,4,14,15^ and homology-independent targeted integration (HITI)^16^, can be utilized for large-scale DNA insertions. However, since these methods rely on generating double-strand breaks (DSBs), indels are also subsequently produced at the target site. Furthermore, DSBs often lead to unwanted large DNA deletions^17,18^, chromosomal aberrations^19^, and cell death^20^, limiting further applications. Thus, several methods have been developed to realize large-scale DNA insertions or deletions without generating DSBs. For example, PE-combined recombinase-based tools, such as PASSIGE^10^ and PASTE^11^, have been developed; however, these techniques require multiple steps involving the installation of a landing pad and the integration of donor DNA at the target site. Additionally, CRISPR-associated transposon (CAST)^13^, bridge RNA-guided recombination^21^, and RNA retrotransposon-based systems^22,23^ have been employed for large-scale DNA insertion in prokaryotes and eukaryotes; however, the overall editing efficiencies were relatively low in human cells. Moreover, no system has yet enabled precise gene replacement, i.e., the simultaneous large-scale deletion and insertion of genes in the genome.

To address these limitations, this study aimed to develop a large-scale DNA replacement tool, named Prime Assembly (PA), which adapts PE tools to produce 3’-flaps of single-stranded DNA (ssDNA) on both genome and donor plasmids. These 3’-flaps anneal to each other precisely, similar to “click chemistry” in small molecules or “Gibson assembly” in DNA oligonucleotides, and initiate strand exchange^5,24^, allowing highly efficient megabase-scale genomic excision and/or multi-kilobase donor insertion at the gene of interest in human cells. We achieved a replacement efficiency of 57.8% at endogenous loci with multi-kilobase payloads using the PA system, highlighting the potential of this platform for therapeutic genome editing. Ultimately, site-specific replacement of large-scale coding sequences (CDSs) in disease-related genes can restore gene function across numerous patients with different mutations, providing a “universal drug” for therapeutic genome editing.

## Results

### Three different strategies for PA

Previously, PE was employed to create a 3’-flap with several nucleotides (nt) at the genome locus for precise duplication of megabase-scale chromosomal DNA in human cells^25^. Subsequently, we hypothesized that creating a 3’-flap at the target genome and a complementary 3’-flap at the donor DNA could induce robust hybridization, ultimately promoting large DNA insertion or replacement. Thus, we proposed three different strategies: (i) generate a single 3’-flap (named A) at the target site and a complementary 3’-flap (named A′) in donor DNA using respective PE guide RNA (pegRNA) to encode a primer binding site (PBS) and a reverse transcriptase template (RTT) for each 3’-flap. We assumed that, after hybridization between the genomic flap (A) and complementary donor flap (A′), the donor-templated DNA extension from the A–A′ hybridization site would be incorporated into the genome through homology-dependent strand invasion. This strategy was named single-flap PA (SF-PA) (**Fig. 1a**); (ii) most processes are similar to the generation of the SF-PA; however, to enhance the invasion of donor homology sequences, a further nick was created by additional nicking guide RNA (ngRNA) on the opposite strand near the homologous sequence in the genome. This strategy was named SF-PA with a nick (SFn-PA) (**Fig. 1b**); (iii) four corresponding pegRNAs were utilized to generate two different 3’-flaps (named A and B) at two target sites, and two complementary 3’-flaps (named A′ and B′) in the donor DNA. For strategy iii, we assumed that after hybridization between genomic flaps and complementary donor flaps (i.e., A–A′ and B–B′), the DNA extensions from each hybridization site would pair with the donor template and be incorporated into the genome, thereby replacing the intervening genomic sequences. This strategy was named dual-flap PA (DF-PA) (**Fig. 1c**).

**Fig. 1.**
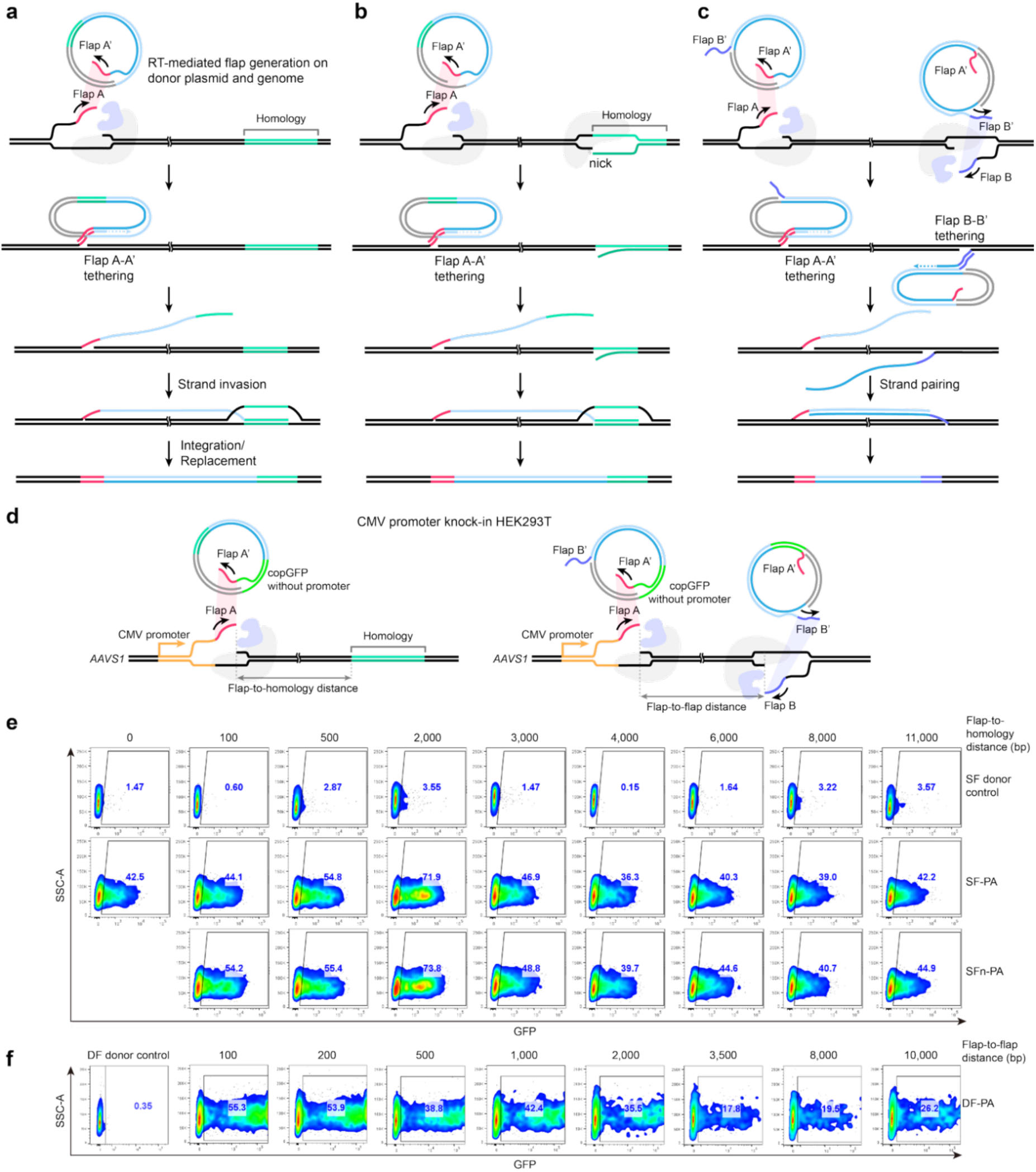
Overview of Prime Assembly. **a–c**. Schematics of single-flap Prime Assembly (PA) (SF-PA), nicked single-flap PA (SFn-PA), and dual-flap PA (DF-PA). Complementary 3′-flaps, generated by pegRNAs, tether the genome and donor to enable donor-templated synthesis and sequence replacement. An additional nick in SFn-PA enhances strand invasion, while paired flaps in DF-PA mediate excision and insertion of large fragments. **d**. Schematic of PA function in HEK293T cells with CMV promoter knock-in at the *AAVS1* locus using promoter-less CopGFP-containing donors; left, SF-PA and SFn-PA; right, DF-PA. **e**. Flow cytometry analysis of integration efficiency of SF-PA and SFn-PA according to flap-to-homology distance (0–11,000 bp), with the corresponding SF donor control included for each distance. X-axis: FITC-A; Y-axis: SSC-A. **f**. Flow cytometry analysis of integration efficiency of DF-PA according to flap-to-flap distance (100–10,000 bp), compared with the DF donor control. X-axis: FITC-A; Y-axis: SSC-A.

To examine these three PA strategies in human cells, we established a cytomegalovirus (CMV) promoter knock-in HEK293T cell line, in which the CMV promoter and blasticidin-S deaminase (BSD) were located in the *AAVS1* safe-harbor region (**Fig. S1**). Then, a donor plasmid encoding copGFP without a promoter was prepared. For the SF-PA and SFn-PA donor plasmids, one flap could be made upstream of copGFP, while the homologous sequences were located downstream of copGFP. In contrast, two different flaps could be made, both upstream and downstream of copGFP for the DF-PA donor plasmid (**Fig. 1d**). When the copGFP cassette (4.3 kbp for SF and SFn; 2.9 kbp for DF-PA) was precisely integrated downstream of the CMV promoter region using the PA system, flow cytometry could be used to detect green fluorescence signals. Initially, the distance between the 30-nucleotide flap and 1 kbp homology arm was designed to be 100 bp for SF-PA and SFn-PA, while the distance between two 30-nucleotide flaps in the genome for DF-PA was intended to be 100 bp. Notably, the PA efficiencies for SF-PA, SFn-PA, and DF-PA using PE7^26^ were very high in the flow cytometry results at 44.4%, 54.2%, and 55.3%, respectively (**Fig. 1e** and **1f**). Moreover, PA events were observed in all cases after increasing the flap-to-homology distance on the genome to approximately 11 kbp for SF-PA and SFn-PA, and the flap-to-flap distance to approximately 10 kbp for DF-PA (**Fig. 1f**), which strongly supports the effectiveness of all three PA strategies as gene replacement tools in human cells.

### Optimal conditions for PA

Despite PA tools possessing high gene replacement efficiencies, we suspected that the frequencies of PA events were potentially overestimated in the flow cytometry results, as cells with a replacement in only one allele exhibited an estimated frequency of 100%. Thus, we conducted digital PCR (dPCR) experiments to quantify the genomic DNA level of PA efficiency. For DF-PA, we revisited the copGFP replacement experiments in the CMV knock-in HEK293T cells, while the PA events were also measured by dPCR, regardless of the flap-to-flap distance (**Fig. 2a**). The dPCR results were highly correlated with the flow cytometry results (R = 0.96); meanwhile, the overall efficiencies by dPCR were expectedly lower than those by flow cytometry (**Fig. 2a**).

**Fig. 2.**
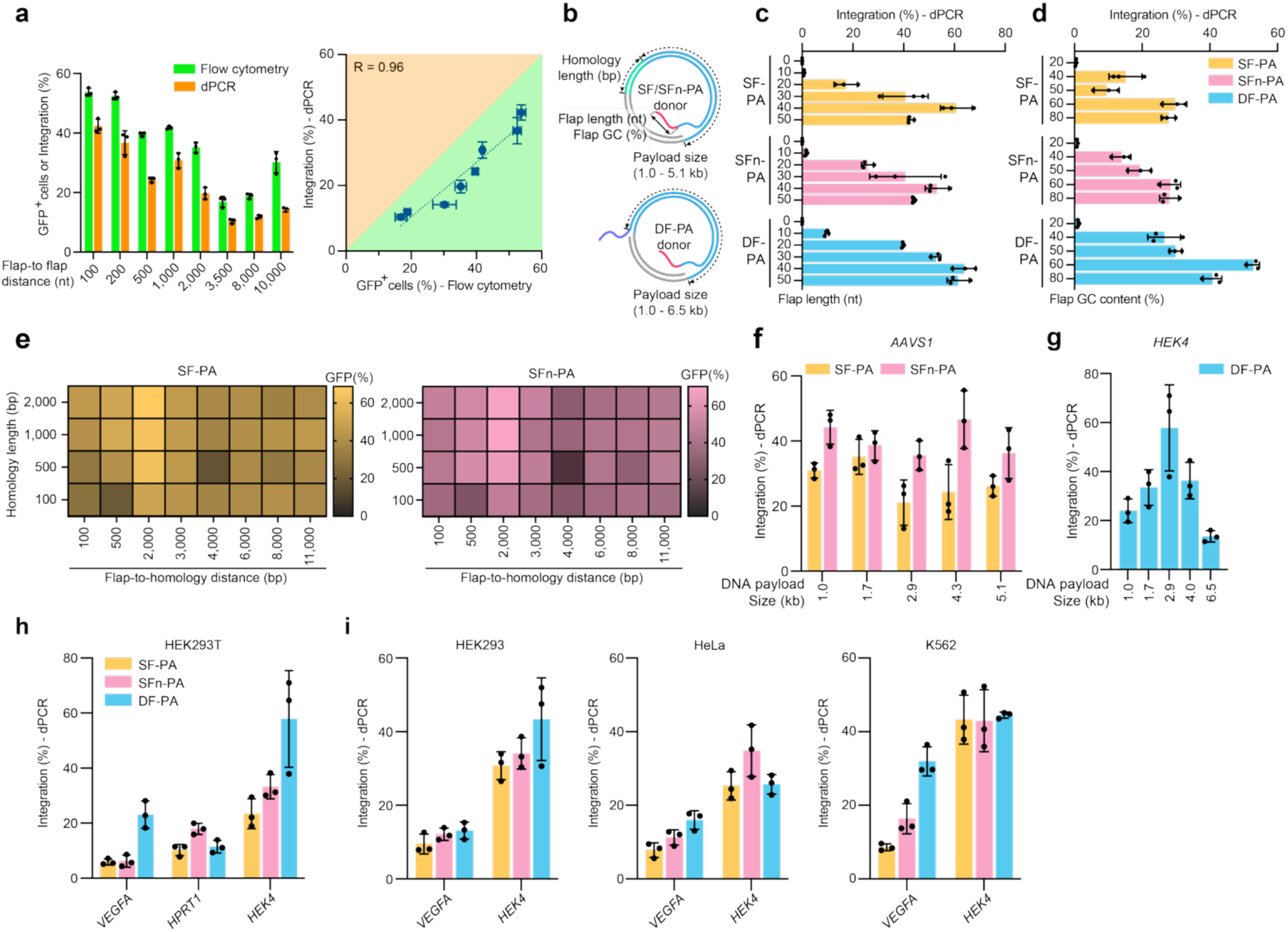
Functional characterization and optimization of PA. **a**. Correlation between DF-PA integration efficiency measured by flow cytometry and dPCR across varying flap-to-flap distances, with a correlation coefficient (R) of 0.96. **b**. Schematic illustrating the optimized elements within donors for each Prime Assembly type. SF-PA and SFn-PA were optimized for homology length (bp), flap length (nt), and flap GC content (%), while DF-PA was optimized for flap length (nt) and flap GC content (%). All donor types, containing promoter-less copGFP sequences, were compared across various payload sizes. **c**. Integration efficiencies of SF-PA, SFn-PA, and DF-PA were measured with flap lengths ranging from 10 to 50 nucleotides (nt). **d**. Integration efficiencies of SF-PA, SFn-PA, and DF-PA were measured using flap GC contents ranging from 20% to 80%. **e**. Integration efficiencies of SF-PA and SFn-PA with varying homology arm lengths and flap-to-homology distances, measured by flow cytometry. **f**. Integration efficiencies of SF-PA and SFn-PA at the *AAVS1* locus with donor payloads of 1.0–5.1 kb. **g**. Integration efficiencies of DF-PA at the *HEK4* locus with donor payloads of 1.0–6.5 kb. **h**. Integration efficiencies of SF-PA, SFn-PA, and DF-PA at multiple endogenous loci (*VEGFA, HPRT1, HEK4*) in HEK293T cells. **i**. Integration efficiencies of SF-PA, SFn-PA, and DF-PA at *VEGFA* and *HEK4* loci in multiple human cell lines (HEK293, HeLa, K562).

Next, we investigated the optimal conditions of the PA tools using flap lengths from 0 to 50 nt and GC contents from 20 to 80% (**Fig. 2b** and **Table S1**). Flap lengths of 30–50 nt and GC contents of 60– 80% promoted efficient replacements with similar trends observed for SF-PA, SFn-PA, and DF-PA (**Fig. 2c** and **2d**). Thus, we selected a standard flap length condition of 30 bp and GC content of 60%. We also examined the optimal homologous arm length (100 bp to 2000 bp) for SF-PA and SFn-PA, with various flap-to-homology distances. Here, the PA efficiency was influenced by both the arm length and flap-to-homology distance, while the highest efficiency was predominantly achieved using arm lengths of 1000–2000 bp (**Fig. 2e**). Therefore, a standard homology arm length condition of 1000 bp was chosen for SF-PA and SFn-PA. Finally, we examined the capacity of payload size for all PA systems, testing payload sizes ranging from 1.0 to 5.1 kbp in the *AAVS1* target for both SF-PA and SFn-PA. Notably, the PA events for each SFn-PA tested payload size and PA efficiency (average 36.2%) were generally higher than those for SF-PA (average 27.6%) (**Fig. 2f**). When an additional nick was induced in the target strand instead of the opposite strand, the PA efficiency for SFn-PA remained consistent over SF-PA (**Fig. S2**), indicating that the additional nick on the opposite strand is critical for enhancing PA efficiency. Next, payloads ranging from 1.0 to 6.5 kbp in size were tested in the *HEK4* target for DF-PA. All cases presented PA events, with a maximum of 57.8% occurring with a 2.9 kb donor (**Fig. 2g**).

To examine whether the PA system could function in different endogenous sites and cell lines, we tested three PA tools in HEK293T cells at the *VEGFA, HPRT1*, and *HEK4* loci, alongside *VEGFA* and *HEK4* in HEK293, HeLa, and K562 cells. The PA efficiencies varied but PA replacement occurred in all endogenous targets with the highest efficiencies noted for *HEK4* in HEK293T cells (average: 23.5%, 33.2%, and 57.8% for SF-PA, SFn-PA, and DF-PA, respectively; **Fig. 2h**). Moreover, we found that PA replacement demonstrated similar PA efficiencies across endogenous targets in all tested cell lines (**Fig. 2i**). These findings indicate that PA tools enable efficient large-scale replacement at multiple endogenous loci across various human cells.

### On-target and off-target analysis of PA outcomes

Oxford Nanopore sequencing was performed in the CMV knock-in HEK293T cell line to validate the accuracy of PA-mediated replacement at on-target sites (4.3 kbp donors for SF-PA/SFn-PA, and a 2.9 kbp donor for DF-PA; **Fig. 3a**). Briefly, genomic DNA was prepared and dephosphorylated. Then, Cas9–ribonucleoprotein (RNP) complexes were treated to cleave upstream of the PA target region, exposing phosphate groups exclusively in the Cas9–RNP cleaving fragments (i.e., on-target site)^27^. These fragments were ligated with unique molecular identifiers (UMIs)-containing adapters, amplified by PCR, and subjected to long-read sequencing. The resulting PA-mediated integration efficiencies for SF-PA, SFn-PA, and DF-PA were 7.3%, 18.6%, and 17.2%, respectively (**Fig. 3b**). Among the integration reads, the exact integration (insertion accuracy) for SF-PA, SFn-PA, and DF-PA was 99.4%, 99.2%, and 96.0%, respectively, with unwanted mutation rates of 0.6%, 0.8%, and 4.0%, respectively (**Fig. 3b**). These data indicate that SF-PA and SFn-PA possess a highly precise integration feature, in contrast with DF-PA.

**Fig. 3.**
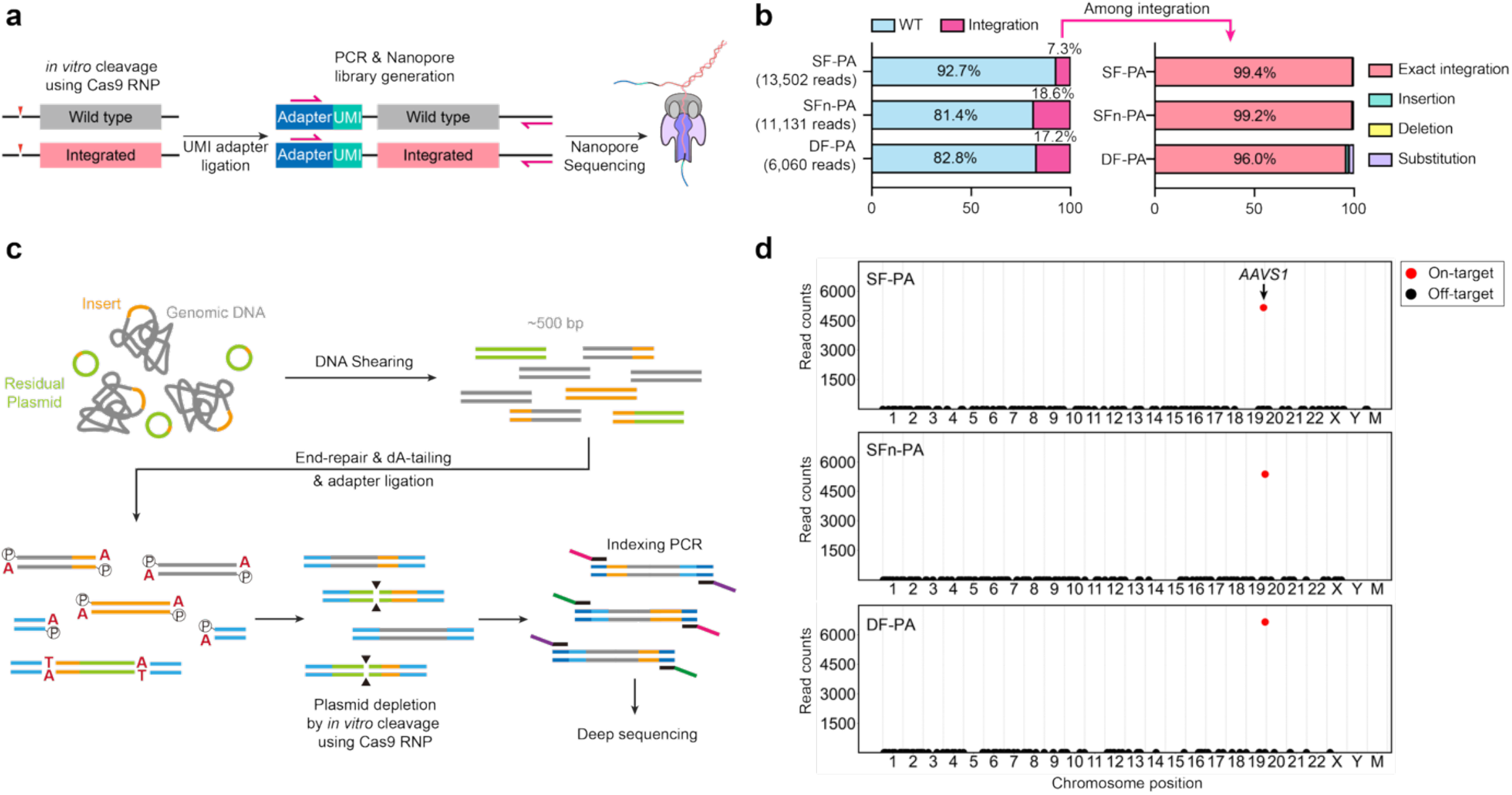
Long-read sequencing validation and off-target assessment of PA. **a**. Schematic of a unique molecular identifier (UMI)-based nanopore sequencing strategy. Genomic DNA was dephosphorylated and cleaved with Cas9 RNPs, followed by ligation of UMI adapters, PCR amplification, and nanopore sequencing to classify integration and wild-type alleles. **b**. Classification of sequencing reads for SF-PA, SFn-PA, and DF-PA into wild-type and integration categories. Integration-positive reads were further subdivided into exact, insertion, deletion, or substitution classes based on sequence accuracy across both ends of the integrated region. **c**. Schematic of genome-wide off-target integration assay. Genomic DNA was sheared into ∼500 bp fragments, followed by end-repair and dA-tailing, then adapter ligation. Residual plasmid sequences were depleted by in vitro Cas9–RNP cleavage. The subsequent fragments were amplified by two-step PCR with indexing in the final step, before next-generation sequencing (NGS) library preparation. **d**. Genome-wide profiling of integration events with on-target and potential off-target reads for each condition.

Next, we evaluated off-target integration events in the same experiments using PA tools. Thus, we designed an integration-searching method to identify genome-wide insertion sites using Illumina sequencing, referencing the GUIDE-seq process^28^. Briefly, genomic DNA was prepared and broken into approximately 500 bp fragments. After performing Cas9–RNP digestion to deplete any residual donor plasmids, these fragments were ligated with adapters and amplified by PCR using a forward primer for ligation sequences and a reverse primer for integration sequences (**Fig. 3c**). The resulting libraries enabled genome-wide profiling of insertion events. No significant off-target integration events exceeding 0.06% of total reads were detected for any PA tool. In contrast, on-target integration at the *AAVS1* locus yielded 5386 reads for SF-PA, 5170 reads for SFn-PA, and 6855 reads for DF-PA (**Fig. 3d**).

### Functional mechanism for PA

We investigated the DNA repair pathways associated with PA-mediated replacement by targeting the *HPRT1* locus in HEK293T cells. Hence, we first tested three representative small-molecule inhibitors: AZD7648 for DNA-PKcs^29^, ART558 for Polθ^30^, and B02 for RAD51^31^. Following chemical treatment, cells were transfected with PA components under the noted culture conditions (**Fig. 4a**). Among them, AZD7648 treatment modestly reduced PA efficiencies for SFn-PA and DF-PA (**Fig. 4b**), while ART558 had minimal effects across all PA formats (**Fig. 4c**). However, B02 treatment dose-dependently decreased PA efficacy, with stronger effects observed for SFn-PA and DF-PA than SF-PA (**Fig. 4d**). These results indicate that RAD51 activity plays a substantial role in PA tools, particularly when extended strand exposure and annealing are required.

**Fig. 4.**
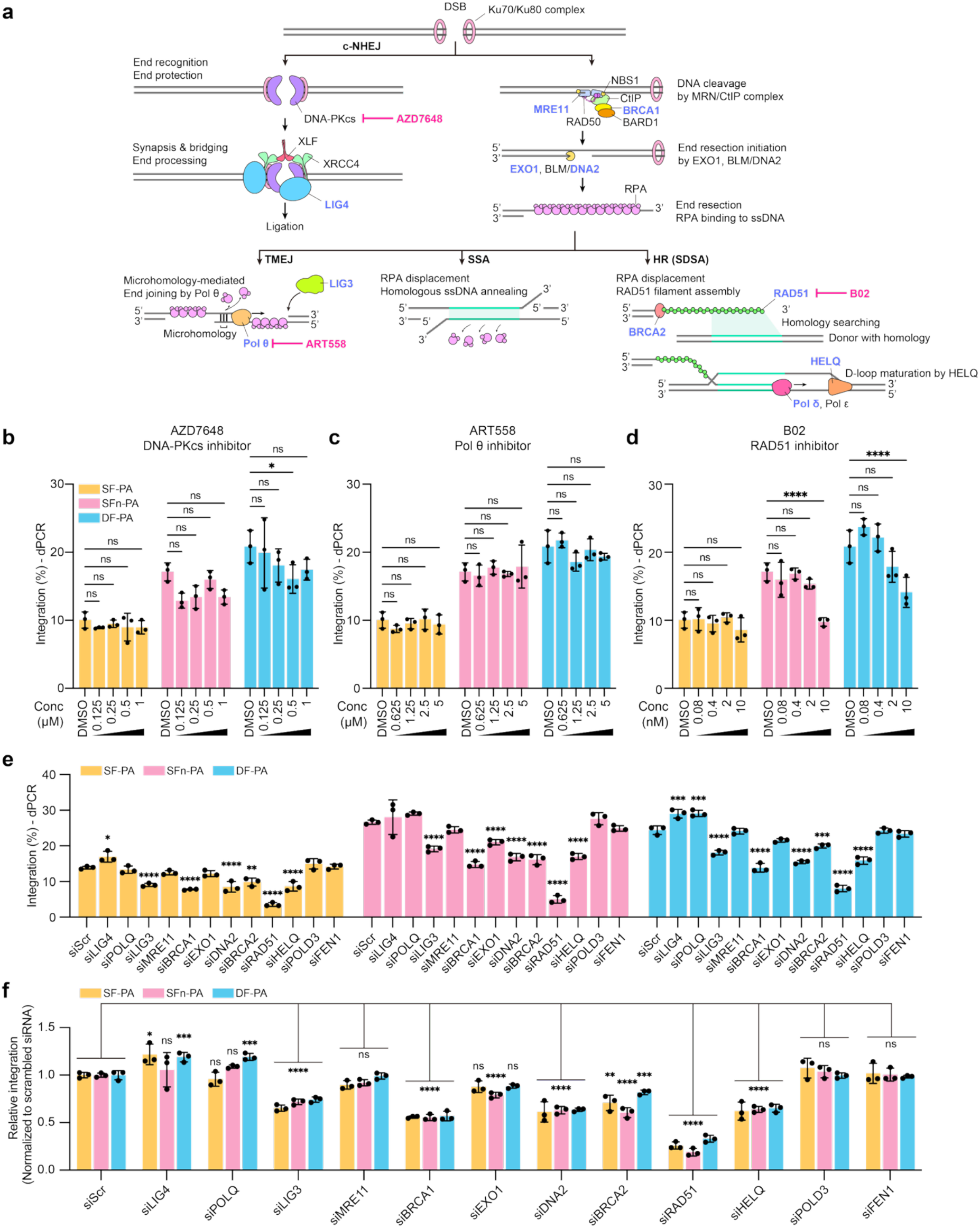
Mechanism of PA. **a**. Schematic of major DNA repair pathways along with their associated factors relevant to PA. Red: factors targeted by small-molecule inhibitors; blue: factors inhibited by siRNA. **b–d**. At the *HPRT1* locus, integration efficiencies of SF-PA, SFn-PA, and DF-PA were measured by digital PCR (dPCR) following treatment with DNA-PKcs inhibitor AZD7648 (b), POLθ inhibitor ART558 (c), or RAD51 inhibitor B02 (d) at the indicated concentrations. **e**. Integration efficiency (%) was measured by dPCR following siRNA-mediated knockdown of the indicated DNA repair genes in SF-PA (yellow), SFn-PA (pink), and DF-PA (blue) cells. Bars represent mean ± SD from three independent experiments (n = 3). Statistical significance was assessed by two-way analysis of variance (ANOVA) comparing each gene-specific siRNA condition to the non-targeting control (siScr) within each PA type. Significance is indicated as follows: ^*^P < 0.05, ^**^P < 0.01, ^***^P < 0.001, ^****^P < 0.0001. **f**. Integration efficiency measured by dPCR in SF-PA (yellow), SFn-PA (pink), and DF-PA (blue) cells after siRNA knockdown of the indicated genes was normalized to the siScr sample within each type. Bars represent mean ± SD from three independent experiments (n = 3). A two-way ANOVA was used to determine statistical significance, with comparisons made to siScr within each PA type. ns = not significant; ^*^P < 0.05, ^**^P < 0.01, ^***^P < 0.001, ^****^P < 0.0001.

To further explore any genetic dependencies, we conducted PA experiments under 13 different siRNA-mediated knockdown conditions (**Fig. 4e**). *LIG3* knockdown, which functions to seal nicks at annealed junctions^32^, reduced PA efficiencies in all three PA formats, most prominently for SFn-PA and DF-PA (**Fig. 4e**). *DNA2* knockdown, which processes 5′-flaps generated during strand-displacement synthesis^33^, promoted a comparable reduction (**Fig. 4e**). Depletion of the homologous recombination (HR) factors, including *RAD51, BRCA1*, and *BRCA2*, significantly reduced the PA efficiency in all three PA formats, with *RAD51* knockdown exerting the most pronounced effect (**Fig. 4e**). *HELQ* depletion also impaired PA efficiency, suggesting a possible role in RAD51/RPA filament remodeling. However, other helicases may also contribute to this process. In contrast, *LIG4* knockdown promoted slight increases for SF-PA and DF-PA, while *POLQ* knockdown increased DF-PA efficiency (**Fig. 4e**). These findings align with the notion that canonical non-homologous end joining (c-NHEJ) and polymerase theta-mediated end joining (TMEJ) can act as competing pathways^34,35^. Normalization to the scrambled siRNA (siScr) control confirmed that all three PA formats operate through broadly similar mechanisms (**Fig. 4f**). These perturbation profiles support a model in which PA utilizes a RAD51-dependent homology-directed mechanism, potentially resembling synthesis-dependent strand annealing (SDSA). Notably, inhibition of mismatch repair (MMR) using PE7 with MLH1-small binder^36^ in HEK293 cells did not significantly alter PA efficiency, indicating that PA operates independently of MMR (**Fig. S3a**).

### Therapeutic applications of PA

Since PA tools are highly efficient for large-scale DNA insertion, deletion, and/or replacement applications, we employed PA therapeutically. Hence, we examined the maximum DNA fragment size that DF-PA could delete and replace within the genome. Thus, endogenous regions spanning 100 kb or 1 Mb were replaced with a donor fragment of 2.9 kb (**Fig. 5a**), yielding efficiencies of 2.5% and 1.9%, respectively, as quantified by dPCR (**Fig. S3b**). PCR amplification using primers positioned approximately 3 kb outside the excision boundaries produced the expectedly sized products. Sanger sequencing confirmed precise junction formation (**Fig. 5b**).

**Fig. 5.**
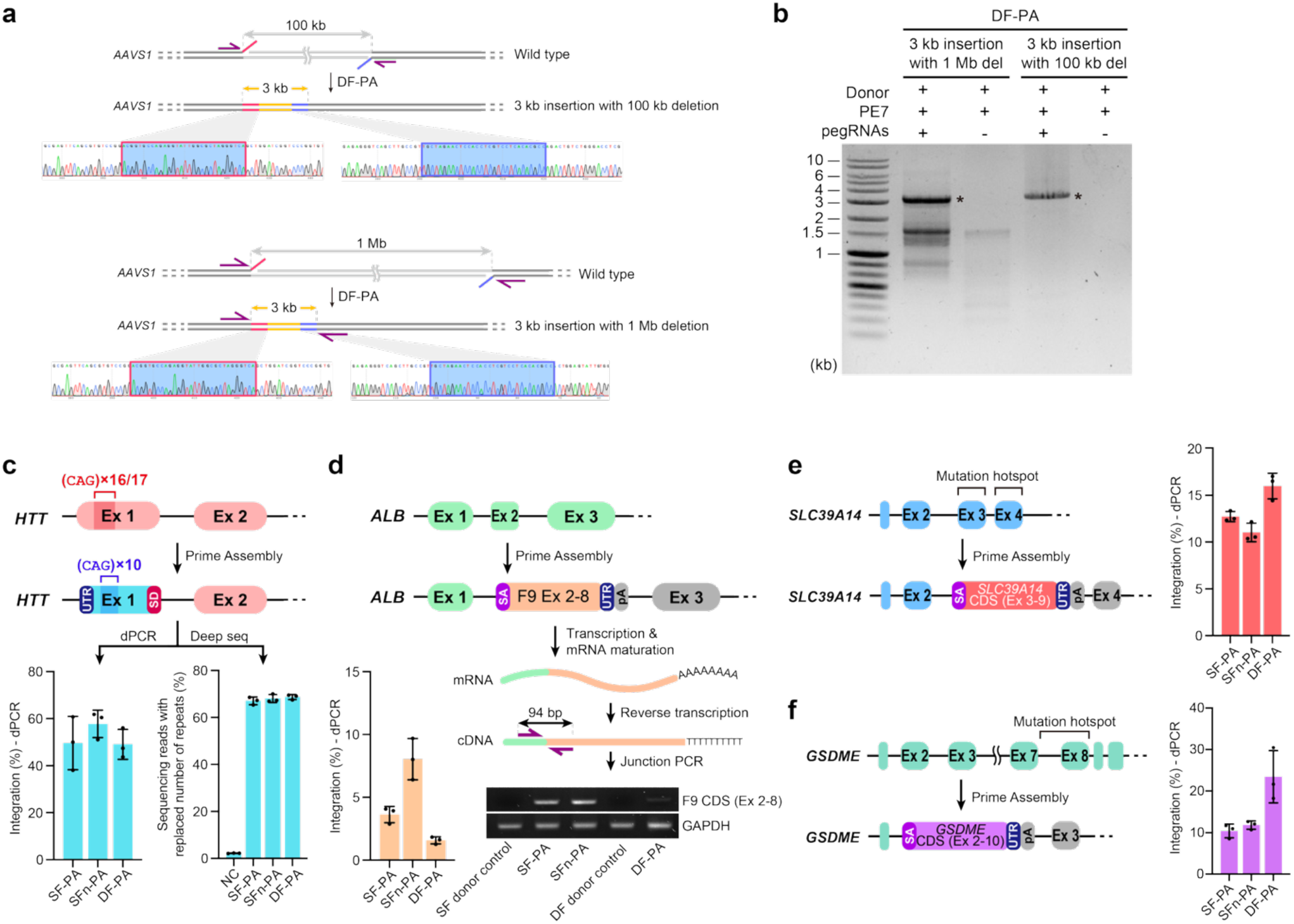
Therapeutic potential of PA. **a**. Megabase-scale replacement at *AAVS1* (DF-PA). Schematic of endogenous segment deletions spanning 100 kb (top) or 1 Mb (bottom) and replacement with a 2.9 kb insert. Long-range PCR amplicons were generated with primers positioned ∼3 kb outside the excision boundaries. **b**. Long-range PCR confirmation. Tris–acetate–EDTA (TAE) agarose gel of DF-PA samples. The expected amplicons for both 1 Mb and 100 kb replacements are clearly detected, appearing only when the donor, PE, and both pegRNAs are transfected. **c**. Exon-level replacement and repeat correction at *HTT*. Designs for SF-PA, SFn-PA, and DF-PA targeting exon 1 (CAG repeat). In HEK293T cells, dPCR confirmed integration for all formats, while deep sequencing (Illumina MiniSeq) verified precise exon 1 replacement with a reduction in the expanded CAG tract. **d**. Liver-specific transgene expression from the *ALB* safe harbor. A splice acceptor–donor carrying *F9* exons 2–8 (hFIX-Padua) was integrated into intron 1 to splice with endogenous exon 1. In HuH7 cells, dPCR detected integration, and cDNA junction PCR yielded the expected ∼94 bp band, indicating proper splicing and expression. **e**. Bypassing a mutation hotspot in *SLC39A14*. To circumvent pathogenic variants across exons 3 and 4, a splice-acceptor donor encoding exons 3–9 was inserted downstream of exon 2. dPCR confirmed integration, restoring a continuous open reading frame. **f**. Correction of splicing-defect alleles at *GSDME*. Donors encoding the full coding sequence (exons 2–10) were integrated into intron 1 to re-establish full-length transcripts that include exon 8. In HEK293T cells, dPCR confirmed integration across formats.

We next applied PA tools to replace STRs in the *HTT* gene, where a CAG trinucleotide expansion in exon 1 promotes the onset of Huntington’s disease^37^. As a proof-of-concept, we attempted to replace exon 1 containing 16 or 17 CAG repeats with synthesized exon 1 containing 10 CAG repeats in HEK293T cells (**Fig. 5c**). PA-mediated replacement efficiencies were measured using dPCR and averaged 49.7% for SF-PA, 57.8% for SFn-PA, and 49.1% for DF-PA (**Fig. 5c**). Illumina sequencing confirmed the precise replacement of exon 1 and correction of the expanded CAG repeat tract, with a reduced repeat copy number observed in 67.09%, 68.08%, and 68.71% of the sequencing reads for SF-PA, SFn-PA, and DF-PA, respectively. These findings highlight the potential of PA to replace entire exons at disease-relevant loci while modulating repeat numbers.

To explore therapeutic transgene integration, we targeted the hepatocyte-specific *ALB* locus, a genomic safe harbor for liver-directed gene therapy (**Fig. 5d**)^38^. A donor construct possessing a splice acceptor (SA) was integrated into intron 1, enabling splicing with endogenous exon 1, which encodes the native secretion signal. As a therapeutic model, we inserted the *F9* fragment (exons 2–8, which encode human factor IX) into patients with hemophilia B. Integration in HuH7 cells reached efficiencies of 3.6% for SF-PA, 8.0% for SFn-PA, and 1.6% for DF-PA (**Fig. 5d**). Junction PCR with complementary DNA (cDNA) revealed an approximately 94 bp band size, confirming the occurrence of proper splicing and expression. This strategy establishes the feasibility of liver-specific expression of therapeutic proteins via PA.

We further applied PA to replace disease-causing mutations in the *SLC39A14* gene (encoding the ZIP14 metal transporter), which causes hypermanganesemia with dystonia syndrome^39^. To bypass a mutation hotspot spanning exons 3–4, we designed a SA–donor encoding exons 3–9 for insertion downstream of exon 2 (**Fig. 5e**). DF-PA achieved an average integration efficiency of 15.1%, whereas SF-PA and SFn-PA showed lower efficiencies of 13.0% and 10.3%, respectively (**Fig. 5e**). Nonetheless, this approach restored a continuous open reading frame, underscoring the therapeutic relevance of PA for transporter-associated disorders.

Finally, we targeted *GSDME* (also known as *DFNA5*), where mutations around introns 7–8 disrupt splicing and promote exon 8 skipping, leading to autosomal dominant non-syndromic hearing loss (ADNSHL) (**Fig. 5f**)^40^. To correct these mutations, we designed donors with the full GSDME (gasdermin E) CDS (exons 2–10) to promote integration immediately downstream of exon 1. This design allows splicing with endogenous exon 1, replacing mutant alleles and restoring full-length transcripts that include exon 8. DF-PA in HEK293T cells achieved an average integration efficiency of 23.5%, compared with 10.4% for SF-PA and 11.8% for SFn-PA, demonstrating the potential of PA for correcting splicing-defect alleles related to ADNSHL (**Fig. 5f**).

## Discussion

Precise replacement of large genomic segments without DSBs remains technically challenging in genome engineering. This study presents three distinct PA strategies that enable the simultaneous site-specific replacement of megabase-scale genomic excision and/or multi-kilobase donor insertion in human cells. Long-read sequencing confirmed the integrity of PA outcomes, showing precise donor incorporation without major alterations. PA-mediated gene replacements at loci including *HTT, ALB, SLC39A14*, and *GSDME* underscore the broad therapeutic potential of this replacement therapy across neurological, hepatic, and metabolic disease contexts, establishing PA as a gene implantation treatment for genome editing or a universal drug for numerous patients with various mutations.

Meanwhile, two groups have posted similar strategies in bioRxiv, differing mainly in donor format. Liu et al. used PCR-derived linear double-stranded DNA (dsDNA), optionally processed to expose 3′-overhangs, and demonstrated the in-cell modular assembly of multiple fragments^41^. Levesque et al. employed synthetic ssDNA and processed 3′-overhang dsDNA for donors. In comparison, our PA strategy utilizes circular dsDNA donors that do not require PCR amplification or in vitro processing, simplifying the workflow while supporting relatively high efficiencies even for multi-kilobase inserts. Within this framework, SF-PA predominantly showed slightly higher junctional precision than DF-PA, as indicated by long-read sequencing. Instead, DF-PA simplifies donor design by eliminating the need for homology arms and supports megabase-scale genomic deletions.

Nonetheless, the PA system relying on dsDNA donors can still promote cytotoxicity in many cell types. To overcome this limitation, circular ssDNA (cssDNA) or RNA-templated donors are necessary to potentially reduce cytotoxicity and innate immune activation while preserving integration efficiency, compared to dsDNA^43^. Additionally, the understanding of the detailed mechanism through which the PA process functions remains limited. CRISPR-based genetic screens should be conducted to clarify which genes or repair pathways are responsible for PA and to identify cellular cofactors that influence its efficiency and accuracy^44^. These directions are expected to broaden the therapeutic applicability of PA and advance the system closer to therapeutic implementation for genome editing.

## Materials and Methods

### Plasmid construction

The pegRNAs and ngRNAs were cloned into the pU6-pegRNA-GG-acceptor plasmid (Addgene, 132777). The vectors were digested using BsaI-HFv2 (New England Biolabs, R3733L), and spacers, scaffold, and extensions were ordered as oligonucleotides synthesized by Macrogen (Seoul, South Korea) and Bionics (Seoul, South Korea), then treated with T4 polynucleotide kinase (Enzynomics, M005S), and annealed (37 °C for 30 min, 95 °C for 2 min, 25 °C for 5 min, with reductions of 0.1 °C/second). The annealed oligos were ligated into the digested vectors using T4 DNA ligase (Enzynomics, M001S) and then transformed into DH5α competent cells, which were prepared using the Mix and Go kit (Zymo Research, T3001). For PA-related donor plasmid construction, cargo sequences were cloned into the pUC19 backbone using Gibson Assembly with the HiFi Assembly Mix (NEB), following the manufacturer’s protocol.

### Mammalian cell culture conditions

HEK293T (ATCC, CRL-11268), HEK293 (ATCC, CRL-1573), HeLa (ATCC, CLL-2), and Huh7 cells (Korean Cell Line Bank, KCLB, Seoul, Republic of Korea) were cultured in Dulbecco’s modified Eagle medium (DMEM; Welgene, LM001-05) supplemented with 10% fetal bovine serum (FBS; Welgene, PK004) and 1% antibiotics (Welgene, LS203-1) at 37 °C with 5% CO_2_. For passaging, the cells were washed with Dulbecco’s phosphate-buffered saline (DPBS; Welgene, LB001-01). Then, the HEK293T and HEK293 cells were detached using 0.05% trypsin–ethylenediaminetetraacetic acid (EDTA; Welgene, LS015), while HeLa and Huh7 cells were detached using 0.25% trypsin–EDTA (Welgene, LS015). The reaction was stopped by adding fresh media (DMEM, 10% FBS, 1% antibiotics), and a portion of the cells was transferred to the next cell culture dish. K562 cells (Korean Cell Line Bank, KCLB, Seoul, Republic of Korea) were cultured in RPMI 1640 medium (Welgene, LM011-01) supplemented with 10% FBS (Welgene, PK004) and 1% antibiotics (Welgene, LS203-1) at 37 °C with 5% CO_2_.

### Construction of CMV promoter–KI cell line

To construct the CMV promoter–KI cell line, HEK293T (ATCC, CRL-11268) cells were electroporated using NxT Electroporation (Invitrogen, N10096). For HDR, 375 ng of p3s–Cas9–HN (Addgene, 104171), 500 ng of pCMV–BSD donor plasmid, and 125 ng of *AAVS1* targeting sgRNA were used (totaling 1000 ng), and cells containing the plasmid were selected at 1 week post-transfection by administering 5 µg/mL blasticidin. After 2 weeks, the selected cells were detached using 0.05% trypsin– EDTA and seeded into 96-well plates (SPL, 30096) at a density of 0.3 cells per well with 100 µL of media (DMEM, 10% FBS, 1% antibiotics) to isolate single clones. After 2 weeks, single clones were harvested, and a portion of the cells was transferred to a new culture plate; the others were lysed in 50 µL of DNA extraction buffer (40 mM Tris, pH 8.9, 1% Tween-20, 0.2 mM EDTA, proteinase K solution (20 mg/mL), 0.2% nonidet P-40). The samples were vortexed for 15 seconds, and then the extracted gDNA was amplified using PCR under the following conditions: 60 °C for 15 minutes, followed by 98 °C for 5 minutes. The PCR products were analyzed by gel electrophoresis on a 1% Tris–borate– EDTA (TBE) agarose gel and subjected to Sanger sequencing. Clones with a single band indicative of the homozygous knock-in were selected and expanded.

### Transfection and genomic DNA extraction

The day before transfection, cells were seeded into 48-well plates (SPL, 30048) at a density of 0.3 × 10^5^ cells per well with 250 µL of medium (DMEM, 10% FBS, 1% antibiotics). The next day, prepared vectors were mixed and treated with 0.65 µL of jetOPTIMUS (Polyplus, 101000006), according to the manufacturer’s protocol. Typically, 300 ng of pCMV-PE7 (Addgene, 214812) and 200 ng of the donor plasmid and 40 ng of two pegRNAs were used for SF-PA transfection, while an additional 40 ng of ngRNA was added for the SFn-PA transfection (a total of 620 ng). Meanwhile, DF-PA transfection predominantly involved 300 ng of pCMV-PE7 (Addgene, 214812), 200 ng of the donor plasmid, and 30 ng of four pegRNAs (totaling 620 ng). After 24 hours, 200 µL of medium was removed and replaced with fresh culture medium (DMEM, 10% FBS, 1% antibiotics). After an additional 48 hours, cells were harvested using 0.05% trypsin–EDTA (Welgene, LS015). The harvested cells were centrifuged at 455 x g for 3 minutes to collect the cell pellet, followed by the addition of 50 µL of DNA extraction buffer (40 mM Tris, pH 8.9, 1% Tween-20, 0.2 mM EDTA, proteinase K solution (20 mg/mL), 0.2% nonidet P-40). The samples were vortexed for 15 seconds and then incubated in a PCR machine under the following conditions: 60 °C for 15 minutes, followed by 98 °C for 5 minutes.

### Electroporation

K562 electroporation was performed using NxT electroporation (Invitrogen, N10096). For SF-PA transfection, 600 ng of pCMV-PE7 (Addgene, 214812) and 400 ng of the donor plasmid, as well as 80 ng of two pegRNAs, were typically used, and an additional 80 ng of ngRNA was added for SFn-PA transfection (totaling 1240 ng). For DF-PA transfection, 600 ng of pCMV-PE7 (Addgene, 214812), 400 ng of the donor plasmid, and 60 ng of four pegRNAs were typically used (totaling 1240 ng). For each reaction, 0.5–1.0 × 10^5^ cells were mixed with plasmids in 10 µL of Neon Resuspension Buffer R (Invitrogen, MPK1096). After electroporation, the cells were cultured in a 37 °C incubator with 5% CO_2_ for 72 hours.

### Chemical inhibition and siRNA knockdown

For chemical inhibition, cells were pretreated with small-molecule inhibitors dissolved in dimethyl sulfoxide (DMSO) or with a vehicle control for 3 h before PA transfection. Media were exchanged 6 h post-transfection and refreshed again at 24 h. Cells were collected for genomic DNA analysis 72 h later.

For siRNA knockdown, cells were transfected with the indicated siRNAs using Lipofectamine RNAiMAX. After 48 h, cells were dissociated, replated at 3.0 × 10^4^ cells per well, and co-transfected with siRNAs and PA plasmids using jetPRIME. Media were exchanged 6 h later and refreshed after 24 h. Cells were harvested for genomic DNA extraction at 72 h post-transfection.

### Electrophoresis and Sanger sequencing

PCR was performed using KOD Multi and Epi (TOYOBO, KME-101) according to the manufacturer’s manual under the following conditions: 98 °C for 2 minutes, 35 cycles of 98 °C for 10 seconds, 56 °C for 20 seconds, and 68 °C for 30 seconds/kb, followed by a final extension at 68 °C for 1 minute. The PCR products were electrophoresed on a 1% Tris–acetate–EDTA (TAE) agarose gel. The bands were gel-extracted using Expin Gel SV (GeneAll Biotechnology, 102-102). The extracted product was analyzed using Sanger sequencing (Cosmogenetech).

### Flow cytometry

At 72 hours post-transfection, cells were harvested using 0.05% trypsin–EDTA (Welgene, LS015). The harvested cells were centrifuged at 455 x g for 3 minutes to collect the cell pellet, and then resuspended in 200 µL flow cytometry buffer (DPBS, 2% FBS). Fluorescence was measured using a BD FACSymphony A5 Cell Analyzer, BD LSRFortessa X-20 instrument, and analyzed using FlowJo v.10 software.

### Digital PCR

The QIAcuity Digital PCR System (Qiagen, Hilden, Germany) was used to quantify genomic integration. One primer pair targeted the integrated junction, while another pair targeted the wild-type sequence to quantify genomic DNA. Each primer set included a probe positioned within the amplicon: a FAM-labeled probe binding to the junction containing the 5′ region of the integrated donor or flap sequence, and (ii) a HEX-labeled probe binding to a separate wild-type sequence serving as an internal reference for normalization. All primers and probes were synthesized by Macrogen (Seoul, South Korea) and Bionics (Seoul, South Korea).

A 12 µL reaction mixture was prepared by combining 0.8 µL of each forward and reverse primer (10 pmol), 0.4 µL of each FAM- and HEX-labeled probe (10 pmol), 3 µL of 4× QIAcuity Probe Master Mix (Qiagen, Hilden, Germany), 4 µL of nuclease-free water, and 1 µL of extracted gDNA. A total reaction mixture of 10.5 µL was loaded into each well of a QIAcuity Nanoplate 8.5k 96-well plate (Qiagen, 250021), followed by application of a sealing cover. PCR cycling conditions were as follows: an initial denaturation at 95 °C for 2 minutes; 40 cycles of 95 °C for 15 seconds, and 55 °C for 30 seconds. Fluorescence was measured using the QIAcuity One 2plex system (Qiagen, 911001) and analyzed with the QIAcuity Software Suite.

### Deep sequencing and data analysis

Illumina sequencers (MiniSeq) were used for genomic data analysis. PCR was performed in two steps for targeted deep sequencing: the first step used 2 µL of extracted gDNA with adapter primers, and the second step used 1 µL of the product from the first step as a template with index primers. PCR was performed using KOD Multi and Epi (TOYOBO, KME-101) according to the manufacturer’s manual under the following conditions: 98 °C for 2 minutes, 29 cycles of 98 °C for 10 seconds, 56 °C for 20 seconds, and 68 °C for 30 seconds, followed by a final extension at 68 °C for 1 minute. The used adaptor and index primers are listed in Table S3. The final PCR products were purified using Expin™ PCR SV (GeneAll Biotechnology, 103-102) and sequenced using the MiniSeq Mid Output kit (Illumina, FC-420-1004).

### High-quality genomic DNA extraction and preparation

For long-read sequencing and the integration assay, gDNA was extracted from cells 72 hours post-transfection using either the DNeasy Blood and Tissue kit (Qiagen, Hilden, Germany) or the AccuPrep Genomic DNA Extraction kit (Bioneer, Daejeon, South Korea) according to the manufacturer’s protocols. DNA concentration was measured using a Qubit fluorometer (Thermo Fisher Scientific, Waltham, MA, USA).

### Library preparation for Oxford Nanopore sequencing

To generate sequencing-ready libraries with defined ligation sites, Cas9–RNP complexes were prepared by mixing 10 pmol of each sgRNA with 12 pmol of Cas9 protein (Enzynomics, Daejeon, South Korea) in 10× rCutsmart buffer (New England Biolabs, B6004S). The mixture was then incubated at room temperature for 30 minutes.

For RNP treatment, 2.5 µg of gDNA was first dephosphorylated using Quick CIP (New England Biolabs, M0525S) at 37 °C for 30 minutes, followed by heat inactivation at 80 °C for 2 minutes. The dephosphorylated gDNA was then incubated with the preassembled RNP complexes at 37 °C for 1 hour. DNA was purified using AMPure XP beads (0.8×, Beckman Coulter, Brea, CA, USA) and eluted in 42 μL nuclease-free water.

Meanwhile, dA-tailing was performed using the NEBNext dA-Tailing Module (New England Biolabs, E6053) according to the manufacturer’s protocol, with a final elution volume of 9 μL nuclease-free water.

UMI-containing adapter oligos were annealed by heating the upper and lower oligos at 95 °C for 3 min, followed by gradual cooling to room temperature. Adapter ligation was performed using 9 μL of dA-tailed DNA, 10 μL Blunt/TA Ligase Master Mix (New England Biolabs, M0367), and 1 μL of a 1:10 diluted adapter, with incubation conducted at room temperature for 1 hour. DNA was purified using AMPure XP beads (0.8×).

Eluted DNA was quantified using a Qubit Fluorometer, and 100 ng was subjected to PCR amplification with KOD Multi and Epi (TOYOBO, KME-101) using primers designed specifically to the adapter sequences and endogenous target regions. Thermal cycling conditions were as follows: initial denaturation at 98 °C for 2 minutes; 38 cycles of 98 °C for 10 seconds, 60 °C for 20 seconds, and 68 °C for 10 minutes; a final extension at 68 °C for 12 minutes. The PCR products were purified with AMPure XP beads (0.8×).

From the purified PCR product, 200 fmol was used for end preparation with the NEBNext Ultra II End Prep enzyme mix and buffer (from NEBNext Ultra II DNA Library Prep kit for Illumina, New England Biolabs, E7647/E7646), following the manufacturer’s protocol. The end-prepped DNA was then processed using the Oxford Nanopore Ligation Sequencing kit (LSK114, Oxford Nanopore Technologies) according to the manufacturer’s instructions for ligation adapter (LA) attachment and sequencing.

### Library preparation for the integration assay using deep sequencing

For comprehensive genome-wide off-target detection, we employed a modified GUIDE-seq protocol adapted for PA. Library preparation was performed using the NEBNext Ultra II DNA Library Prep kit for Illumina (New England Biolabs, Ipswich, MA, USA).

A total of 1 µg of gDNA was diluted in 130 μL using 1× Tris–EDTA (TE) buffer. DNA fragmentation was performed using a Covaris S2 sonicator (Covaris, Woburn, MA, USA) to yield an average fragment size of 500 bp. Fragmented DNA was purified using SPRI beads (0.7×, Beckman Coulter), washed twice with 80% ethanol, and eluted in 50 μL TE buffer.

End repair was performed using 7 μL of NEBNext Ultra II End Prep Reaction Buffer (New England Biolabs, E7647) and 3 μL of NEBNext Ultra II End Prep Enzyme Mix (New England Biolabs, E7646) with fragmented DNA. The reaction was incubated at 20 °C for 30 minutes, followed by incubation at 65 °C for 30 minutes, and then held at 4 °C.

Custom adapters containing PA-specific sequences were annealed by heating a mixture of upper and lower oligos at 95 °C for 3 minutes, followed by gradual cooling to room temperature to allow duplex formation. Adapter ligation was performed using 60 μL of end-repaired DNA, 2.5 μL of annealed adapters, 30 μL NEBNext Ultra II Ligation Master Mix (New England Biolabs, E7648), and 1 μL NEBNext Ligation Enhancer (New England Biolabs, E7374) to a total volume of 93.5 μL. The reaction was incubated at 20 °C for 15 minutes with the heated lid off, followed by purification using SPRI beads (0.7×) and elution in 25 μL nuclease-free water.

To eliminate residual plasmid DNA containing flap sequences, plasmid-specific cleavage was performed using CRISPR–Cas9–RNP complexes before PCR amplification. Four sgRNAs were synthesized via in vitro transcription: (i) two targeting SpCas9 scaffold sequences (10 pmol each), and two targeting plasmid-specific sequences within the pegRNA downstream extension (10 pmol each). RNP complexes were prepared by mixing the four sgRNAs with 24 pmol of Cas9 protein (Enzymomics, Daejeon, South Korea) in 10× rCutsmart buffer (New England Biolabs, B6004S) with nuclease-free water, to a total volume of 10 μL, and incubated at room temperature for 30 minutes. The digestion reaction was performed by mixing 25 μL of purified ligated DNA with 10 μL of RNP complexes, along with rCutSmart buffer and nuclease-free water, to a total volume of 40 μL. The reaction was incubated at 37 °C for 1 hour, followed by heat inactivation at 72 °C for 5 minutes. DNA was purified using SPRI beads (0.7×) and eluted in 10 μL nuclease-free water.

Primary amplification was performed with PA-specific enrichment primers that target adapter and extension sequences. The PCR mixture contained 12.5 μL of NEBNext Ultra II Q5 Master Mix (New England Biolabs, E7649), 2.5 μL each of forward and reverse enrichment primers (10 μM), and plasmid-depleted genomic DNA, in a total volume of 25 μL. Thermal cycling conditions included an initial denaturation at 98 °C for 30 seconds, followed by 20 cycles of two-step PCR (98 °C for 10 seconds, 72 °C for 30 seconds). PCR products were purified using SPRI beads (0.7×) and eluted in 12 μL of nuclease-free water.

Secondary amplification for next-generation sequencing (NGS) library preparation was performed using indexed primers (D501 and D701). The reaction mixture consisted of 10 μL NEBNext Ultra II Q5 Master Mix (New England Biolabs, E7649), 0.5 μL each of indexed primers (10 μM), and 8.5 μL of primary PCR product, for a total volume of 20 μL. Thermal cycling consisted of 98 °C for 30 seconds, followed by 10 cycles of 98 °C for 10 seconds, 65 °C for 20 seconds, and 72 °C for 15 seconds. Final libraries were purified using SPRI beads (1×), quantified with a Qubit fluorometer, and subjected to deep sequencing.

### Bioinformatics analysis of integration assay

Raw sequencing reads were first filtered to retain only meaningful reads containing specific indicator sequences using custom Python scripts. Reads containing the adapter sequence CTACAAGAGCGGTGAGT in R1 and extension sequence GGTGCCAGAGGTATTGGCGCTAGGGTCA in R2 were selected for downstream analysis.

UMIs were extracted from the filtered reads using UMI-tools (v1.1.2) with a nine-nucleotide barcode pattern (NNNNNNNNN). The UMI-tagged reads were then aligned to both on-target reference sequences and the human reference genome (GRCh38) using minimap2 (v2.24) with parameters optimized for short-read alignment (-ax sr --eqx). Alignment files were converted to the BAM format and indexed using SAMtools (v1.15).

To ensure high-quality alignments, only properly paired and mapped reads were retained following SAMtools filtering (flags -f 0x2 -F 0xC). UMI-based deduplication was performed using UMI-tools dedup function with paired-end mode to remove PCR duplicates while preserving biological diversity.

Primer-derived off-target artifacts were removed using custom filtering criteria based on CIGAR string analysis. Specifically, reads with flag 147 (second in pair, reverse strand) that had right soft-clipping ≤26 nucleotides or reads with flag 163 (second in pair, forward strand) that had left soft-clipping ≤26 nucleotides were excluded, as these likely represented primer-binding artifacts rather than genuine integration events.

Final integration site quantification was performed by analyzing R2 reads and determining the 5’ mapping positions. For forward strand alignments, the leftmost mapping position (reference_start + 1) was recorded, while for reverse strand alignments, the rightmost mapping position (reference_end) was used. Integration sites were counted and tabulated across all chromosomes to identify potential off-target locations.

## Acknowledgements

We thank Dr. James W. Wilson (EssayReview) for English-language editing. We also thank Profs. Jangsup Moon, Seungbok Lee, and Sang-Yeon Lee at Seoul National University Hospital for their helpful discussion. Most of the sequencing data analysis was conducted using the computing server at the Genomic Medicine Institute Research Service Center. This research was supported by grants from the National Research Foundation of Korea (NRF) (No. 2021M3A9H3015389, No. RS-2024-00451880, No. RS-2024-00455559, and SRC-NRF2022R1A5A102641311) awarded to S.B. Additional support was provided by the Korean Fund for Regenerative Medicine (KFRM) (No. RS-2024-00332601), a grant from the Ministry of Food and Drug Safety (No. 25202MFDS003) in 2025, the SNUH Lee Kun-hee Child Cancer & Rare Disease Project (No. 25B-001-0700), and Samsung Research Funding & Incubation Center of Samsung Electronics under Project (No. SRFC-MA2101-06) also awarded to S.B.

## Author contributions

H.J. and S.B. conceived the project; H.J., B.-D.J., and Y.-W.K. cloned plasmids, designed cell experiments, and analyzed data; C.-J.J. and H.U. conducted bioinformatics analysis; H.K. designed and performed the long-read sequencing; Y.E.O. and Y.P. performed cell experiments. S.B. supervised the project. H.J. and S.B. wrote the manuscript with input from all authors.

## Additional information

Supplementary Information accompanying this paper is available at

## Declaration of generative AI and AI-assisted technologies

The authors did not use any AI or AI-assisted technologies during the preparation of this work.

## Code availability

The code related to this study is available at https://github.com/BaeLab/PrimeAssemblyAnalysis

