## Supplementary Information for "Site-specific replacement of large-scale DNA fragments in human cells"

**List of Supplementary Materials:**

Supplementary Figure 1 to 4.

Supplementary Table 1

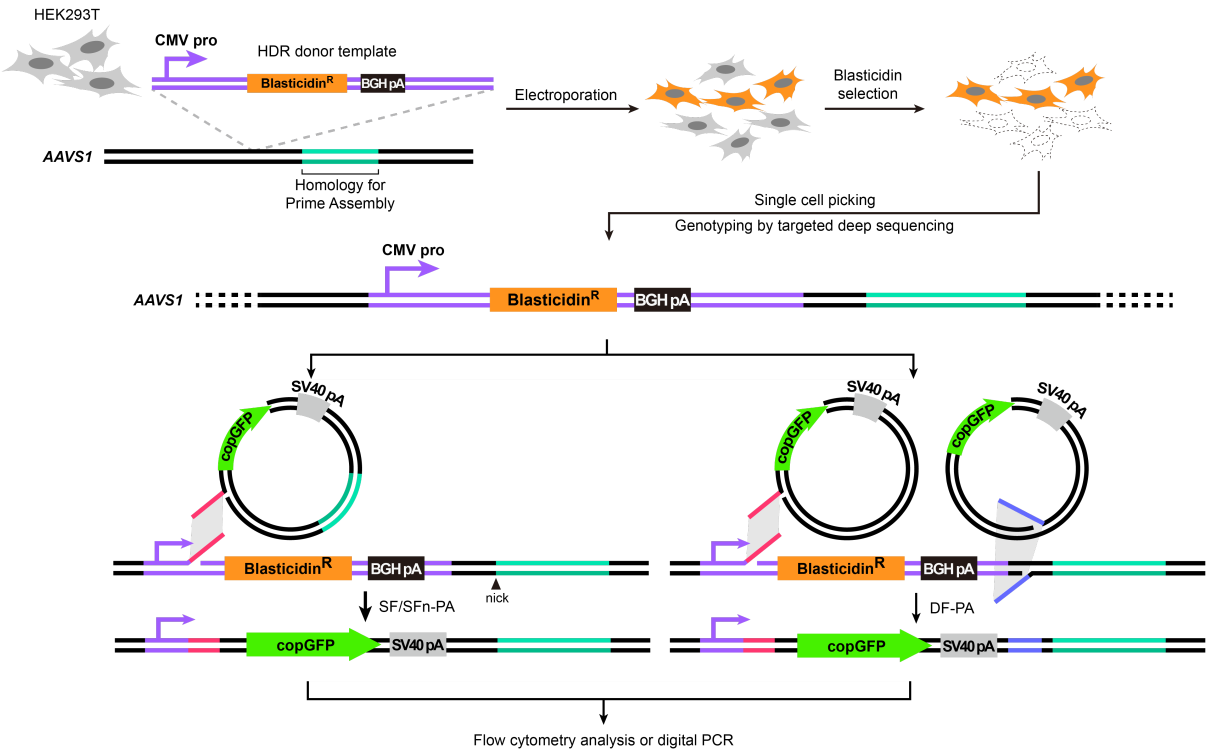


**Supplementary Fig. 1. Generation of CMV promoter knock-in cell line for reporter assay.** Schematic of CMV promoter knock-in cell line generation. A donor containing a blasticidin cassette with the CMV promoter was designed for homology-directed repair targeting the *AAVS1* locus in HEK293T cells. Cas9, sgRNA, and donor DNA were introduced via electroporation. Following blasticidin selection, single cells were picked and genotyped to establish clonal cell lines with successful CMV promoter knock-in. PA was then performed using a promoter-less copGFP donor to replace the blasticidin cassette, allowing integration to be detected by GFP-positive signal in a reporter assay.

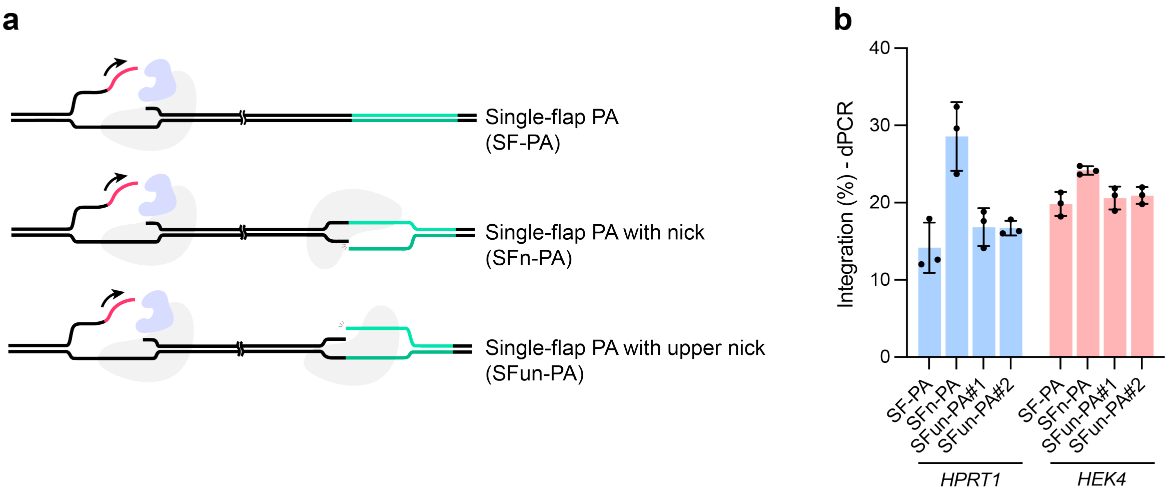


**Supplementary Fig. 2. Evaluation of upper nicking guide in SFn-PA. a.** Schematics of SF-PA, SFn-PA, and SFn-PA with an upper nicking gRNA (upper-ngRNA) that nicks the opposite/top strand as drawn. **b.** dPCR-measured integration in HEK293T cells at *HPRT1* and *HEK4*. Introducing the upper-ngRNA shows little to no improvement over standard SFn-PA.

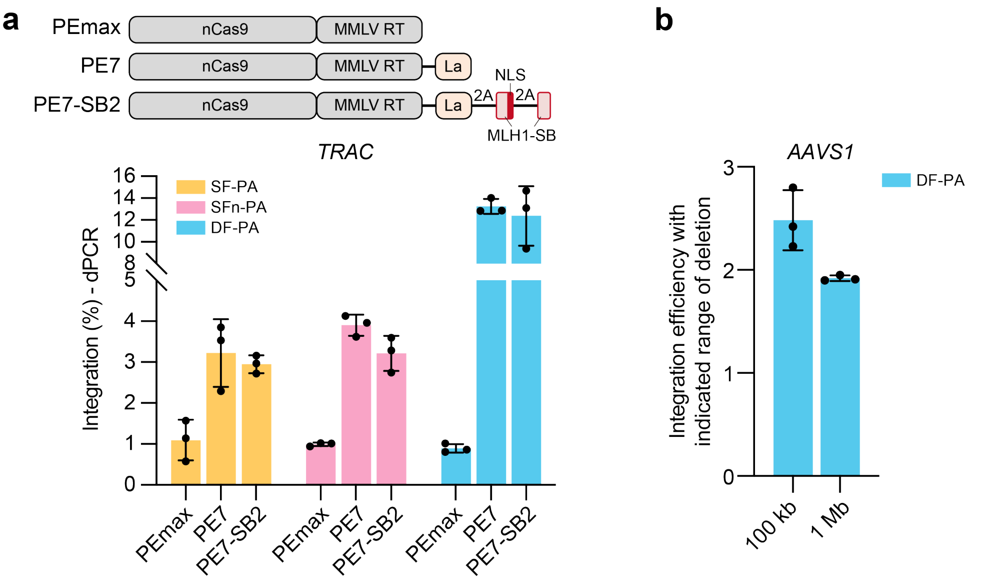


**Supplementary Fig. 3. Comparison of prime editor variants and evaluation of DF-PA in large-scale deletions. a.** Integration efficiencies of SF-PA, SFn-PA, and DF-PA at *TRAC* using three versions of prime editors (PEmax, PE7-SB, PE7-SB2), quantified by dPCR. **b.** Integration efficiencies of DF-PA at *AAVS1*, with largescale deletions of 100 kbp and 1 Mbp, quantified by dPCR.

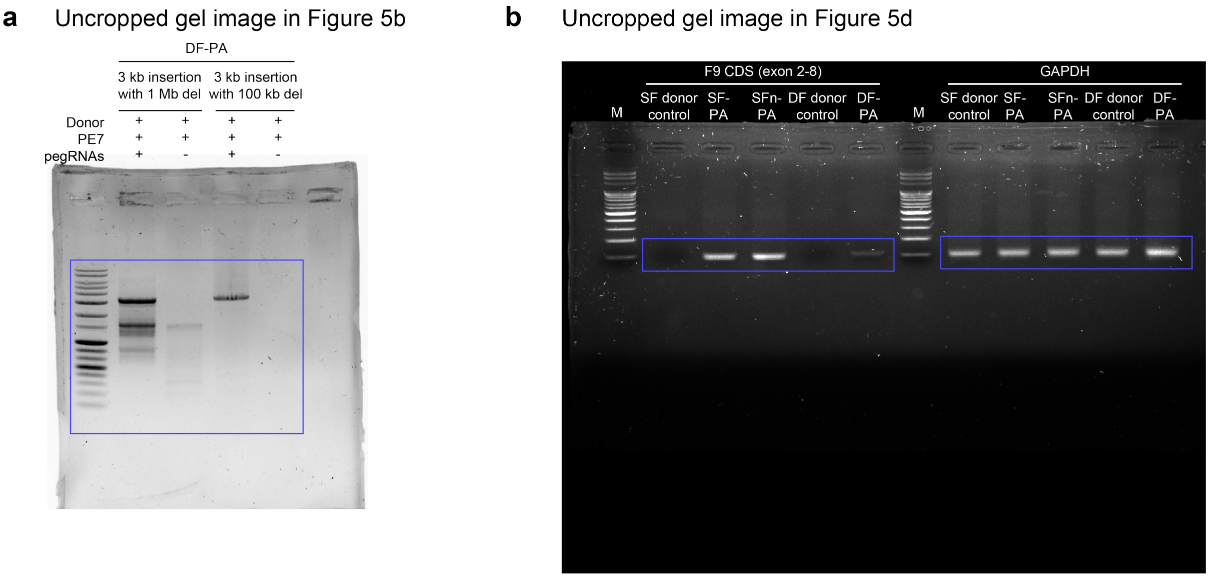


**Supplementary Fig. 4. Uncropped gel images for main figures. a.** Uncropped gel for **Fig. 5b** (long-range PCR confirming 2.9 kb insertion with 100 kb or 1 Mb deletions under DF-PA); cropped regions are boxed. **b.** Uncropped gel for **Fig. 5d** showing cDNA junction PCR for F9 CDS (exons 2–8) and *GAPDH* control; boxed areas correspond to the main figure.

| Flap length optimization | |
|---|---|
| **Flap** | **Sequence (5' to 3')** |
| Flap A (10nt) | TTCGCGAGAA |
| Flap A (20nt) | GCCAATACCTCTGGCACCGT |
| Flap A (30nt) | TGACCCTAGCGCCAATACCTCTGGCACCGT |
| Flap A (40nt) | AGGTGTACCCTGACCCTAGCGCCAATACCTCTGGCACCGT |
| Flap A (50nt) | GCCAGCGTTAAGGTGTACCCTGACCCTAGCGCCAATACCTCTGGCACCGT |
| Flap B (10nt) | CAGCTGATCA |
| Flap B (20nt) | TGCTAGAGCTCCACCTCGTC |
| Flap B (30nt) | TGCTAGAACTCCACCTCGTCCTCACACGCC |
| Flap B (40nt) | TGCTAGAACTCCACCTCGTCCTCACACGCCACTAGGCTCG |
| Flap B (50nt) | TGCTAGAACTCCACCTCGTCCTCACACGCCACTAGGCTCGCCAACGGGTT |
| **Flap GC content optimization** | |
| **Flap** | **Sequence (5' to 3')** |
| Flap A (GC 20%) | ATATTTAATGAAGTTGTAAAATGTTTGGTA |
| Flap A (GC 40%) | GTTGGAGTTATGAATTGTCACACTCATGGA |
| Flap A (GC 50%) | TCAGTAACCCGGAGCTCTAATTCTCgCGAA |
| Flap A (GC 60%) | TGACCCTAGCGCCAATACCTCTGGCACCGT |
| Flap A (GC 80%) | CCGCGTGGCCTGGAGGACCCGTCGGGCCAG |
| Flap B (GC 20%) | TATCTATTATAAGTATAAACACAACTTGAT |
| Flap B (GC 40%) | TCGTACTTGCAACGCTATCATGTCTAGTAA |
| Flap B (GC 50%) | GAAGCGATAGTCGATCGTAGCAGCTGATCA |
| Flap B (GC 60%) | TGCTAGAACTCCACCTCGTCCTCACACGCC |
| Flap B (GC 80%) | CGGAGCCGAGCTGGCGCCTCGCGCAGCGGA |

**Supplementary Table 1. Flap sequences used for optimization of PA, related to Figure 2b–2d.** Flap length optimization (top) and flap GC content optimization (bottom) were systematically tested using the indicated sequences. Sequences highlighted in yellow represent the optimal flap designs that yielded the highest efficiencies under the respective conditions.
